# Automated Structural Variant Verification in Human Genomes using Single-Molecule Electronic DNA Mapping

**DOI:** 10.1101/140699

**Authors:** Michael D. Kaiser, Jennifer R. Davis, Boris S. Grinberg, John S. Oliver, Jay M. Sage, Leah Seward, Barrett Bready

## Abstract

The importance of structural variation in human disease and the difficulty of detecting structural variants larger than 50 base pairs has led to the development of several long-read sequencing technologies and optical mapping platforms. Frequently, multiple technologies and ad hoc methods are required to obtain a consensus regarding the location, size and nature of a structural variant, with no approach able to reliably bridge the gap of variant sizes between the domain of short-read approaches and the largest rearrangements observed with optical mapping.

To address this unmet need, we have developed a new software package, *SV-Verify™*, which utilizes data collected with the Nabsys High Definition Mapping *(HD-Mapping™*) system, to perform hypothesis-based verification of putative deletions. We demonstrate that whole genome maps, constructed from electronic detection of tagged DNA, hundreds of kilobases in length, can be used effectively to facilitate calling of structural variants ranging in size from 300 base pairs to hundreds of kilobase pairs. *SV-Verify* implements hypothesis-based verification of putative structural variants using a set of support vector machines and is capable of concurrently testing several thousand independent hypotheses. We describe support vector machine training, utilizing a well-characterized human genome, and application of the resulting classifiers to another human genome, demonstrating high sensitivity and specificity for deletions ≥300 base pairs.

## INTRODUCTION

Over the past decade, much progress has been made in understanding human genome variation and its implications in human disease through the use of next-generation sequencing (NGS) technologies.^1,2^ While NGS technologies are proficient at the detection of single-nucleotide polymorphisms and indels, the inherent short read length of NGS has proven to be a substantial impediment in the detection of larger variations.^3,4^ Structural variants (SVs) encompass genomic rearrangements involving greater than 50 bp and include deletions, insertions, inversions, and translocations.^5^ Numerous computational methods using NGS data have been developed that evaluate abnormal alignments, discordant read pairs, and changes in coverage to detect SVs.^6^ However, short read length and coverage bias significantly reduce the sensitivity and specificity of SV detection from NGS data. The importance of structural variation in human disease and the limitations of NGS sequencing technologies motivated the development of third-generation sequencing technologies that provide longer read lengths.^7–12^

While third-generation technologies have greatly improved the detection of SVs, bioinformatic bottlenecks remain due to limited read length and high error rates. More recently, DNA mapping of single molecule fragments, in excess of 100 kbp in length, has emerged as an orthogonal approach for the detection of structural variation.^13–16^ Importantly, mapping technologies do not have the same biases as sequencing technologies and can be used in combination with sequencing data to create and refine chromosomal-scale assemblies. Two optical mapping approaches have been released, the Argus system from Opgen and the Irys and Saphyr systems from Bionano Genomics, and have been used to improve genome assemblies.^14,16^ However, even under optimal conditions, the resolution of optical methods is limited by diffraction to approximately 1 kbp (single molecule) and intervals between 1 kbp and 2 kbp are widely under-represented.^17–19^ The resolution limit of optical genome maps limits the minimum size of SVs that can be reliably detected to a size that is significantly larger than the size of SVs that are efficiently detected using NGS methods.

To address the need for long-range structural information while maintaining resolution to complement NGS technologies, Nabsys has developed the *HD-Mapping™* system, which utilizes a completely electronic detection scheme to generate high-density map information from single molecules that are hundreds of kbp in length. Long-range information is preserved so that structural rearrangements, deletions, and duplications are easily identified. Electronic detection and mapping of single DNA molecules by the Nabsys platform has been described.^20^ Briefly, long DNA molecules (ranging from 30 kbp to hundreds of kbp in length) are tagged in a sequence selective manner utilizing commercially available nicking enzymes and a proprietary marker. Tagged DNA molecules translocate through solid-state nanodetectors at a speed exceeding 1 Mbp/sec as the tags on the DNA backbone are electronically detected. The time between consecutive tags is recorded during data collection and the physical distance between tags is accurately determined by accounting for the variation in intramolecular velocity during translocation. Single-molecule reads may be compared to a provided reference sequence using Nabsys proprietary software and a consensus genome map is generated from the data. Single-molecule data from the Nabsys platform have false-positive and false-negative rates below 5% and intervals as small as 300 bp are resolved on single molecules. These errors are stochastic and are rapidly eliminated from consensus maps at very low coverage.^20^ The advantages of electronic sensing include higher resolution, accuracy, and information density of the resulting data versus those provided by optical technologies. In addition the platform is highly scalable and low cost.

We demonstrate that whole genome maps constructed from electronic detection of long DNA can be used to call SVs with high sensitivity and specificity across a wide range of variant sizes. Single-molecule events from the Nabsys *HD-Mapping* platform are collected for the genome of interest. The data is mapped against the original source (SRC) reference plus an aggregate alternate (ALT) reference, representing up to several thousand independent hypotheses. Here, we describe the development of Nabsys *SV-Verify™*, a software package composed of several support vector machines (SVMs) that provides an efficient and robust pipeline for the systematic and automated evaluation of putative SVs. We discuss *SV-Verify* training using reference material from the National Institute of Standards and Technology (NIST) Genome in a Bottle (GIAB) project, for a well-characterized human genome (NA12878).^21,22^ We also report feature selection, SVM kernel selection, training and cross-validation methods. We show the resulting sensitivity and specificity of each SVM and present results obtained by applying *SV-Verify* to several thousand putative deletions within another human genome, NA24385.

## MATERIALS AND METHODS

### DNA Sample Selection

Training of the SVMs contained in *SV-Verify* was performed with data collected from human genome NA12878 (NIST ID: HG001). This genome was extensively characterized by the GIAB project, a public-private-academic consortium hosted by NIST. Deletion calls were made, relative to reference GRCh37, by a large number of consortium members including different technologies and library preparation methods, as well as a variety of tool sets.^22^ *SV-Verify* performance was evaluated with data from a different human genome, NA24385 (NIST ID: HG002) and a list of putative deletions provided by the GIAB consortium. The results presented here were obtained using the v0.1.7 putative deletion set (June 2016) which included calls made by multiple sequencing technologies (Illumina, PacBio, Complete Genomics, and Bionano Genomics) (ftp://ftp-trace.ncbi.nlm.nih.gov/giab/ftp/data/AshkenazimTrio/analysis/NIST DraftIntegratedDeletionsgt19bp v0.1.7).

### Sample Preparation and Data Collection

Human cell lines NA12878 and NA24385 were obtained from Coriell Cell Repositories (Camden, NJ) and were routinely cultured in Roswell Park Memorial Institute 1640 Medium supplemented with 15% fetal bovine serum and 1X antibiotic - antimycotic solution (Life Technologies, Carlsbad, CA) at 37°C and 5% CO2. Human genomic DNA was isolated using the Macherey-Nagel NucleoBond AXG 500 column system (Bethlehem, PA) in conjunction with Macherey-Nagel

NucleoBond Buffer Set IV. DNA samples were nicked with Nt.BspQI (4.4 U/*μ*g) (New England Biolabs, Ipswich, MA) and tagged with Nabsys proprietary chemistry. Samples were then coated with RecA (Enzymatics, Beverley, MA) protein in the presence of ATPγS (Sigma Aldrich, St. Louis, MO). Data were collected utilizing the Nabsys *HD-Mapping* platform and mapped using Nabsys software to the GRCh37 reference. Average coverage depths of 99- and 65-fold were collected for NA12878 and NA24385 respectively.

### Hypothesis-based Deletion Verification

*SV-Verify* employs a hypothesis-based verification strategy to determine the likelihood that a putative deletion exists. First, an ALT reference segment for each putative deletion is generated from the source reference to match the putative deletion hypothesis. Examples of ALT reference segments representing putative deletions in which probe sites are either removed or the spacing between two probes is shortened to create a smaller interval are shown in Figure 1. Flanking intervals on either side of the putative deletion are included in each ALT reference segment to provide sufficient context for unique mapping. Several thousand ALT reference segments may be combined into an aggregate ALT reference to facilitate concurrent testing of thousands of hypotheses. Molecules are allowed to map to either the SRC or ALT reference according to which they most closely resemble.

**Figure 1.**
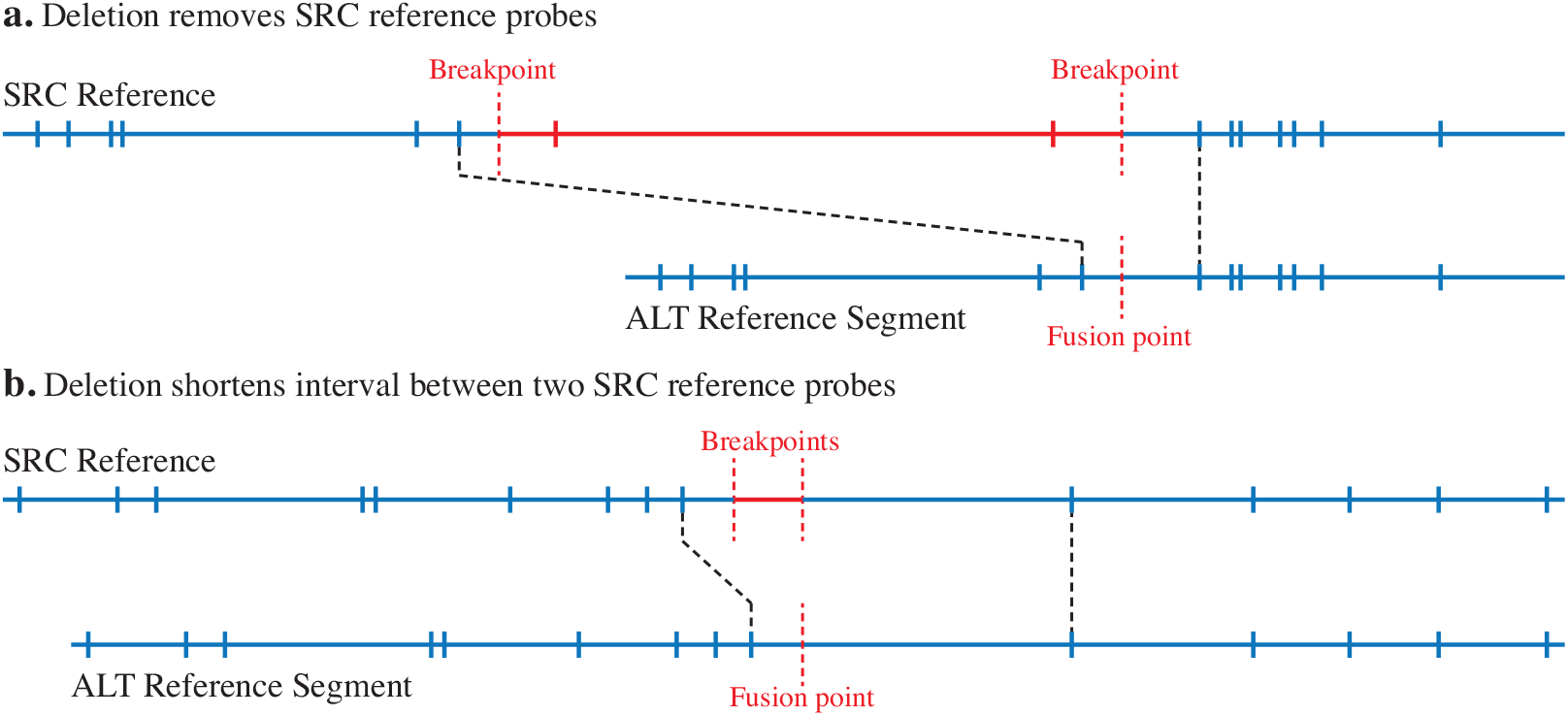
Construction of an ALT reference segment for a putative deletion. Hypothesized deletions resulting in a) removed probes or b) a shortened interval in the ALT reference segment.

When the SRC and ALT reference segment are unique and sufficiently different from each other, the presence and zygosity of a putative deletion can be determined by summing the number of mapped molecules sufficiently spanning the breakpoints of the SRC reference and the number of mapped molecules sufficiently spanning the corresponding fusion point of the ALT reference segment. There are several possible outcomes: 1) Deletion not present: many reads span the SRC breakpoints and few reads span the ALT fusion point. 2) Heterozygous deletion: approximately equal number of reads span the SRC breakpoints and the ALT fusion point. 3) Homozygous deletion: few reads span the SRC breakpoints and many reads span the ALT fusion point. 4) No call: insufficient number of reads span either the SRC breakpoints or ALT fusion point.

It is important to note that although a putative deletion is asserted with a reference start location and size, the hypothesis being tested is that the specified deletion size has occurred between a pair of reference probe locations, potentially deleting additional reference probes (Figure 1). If another deletion or insertion exists within the same region, we may refute the asserted hypothesis. To increase the utility of *SV-Verify* calls, for each putative deletion, output files contain chromosome, putative start, putative size, and SVM score, as well as the location of the two reference probes bounding the putative deletion.

A screenshot of our analysis software demonstrating mapping of single-molecule reads to the SRC reference and the corresponding ALT reference segment for a 43,364 bp homozygous deletion in human genome NA12878 is shown in Figure 2. The upper panel is a software screen image showing two breakpoints for a putative deletion (dashed red lines) on the SRC reference. Reads do not map in the region between the breakpoints on the SRC. The lower panel shows the ALT reference segment constructed by removing the section corresponding to the putative deletion from the SRC reference. Many reads map across the fusion point in the ALT reference segment. This pattern of mapped reads on SRC and ALT references indicates a homozygous deletion.

**Figure 2.**
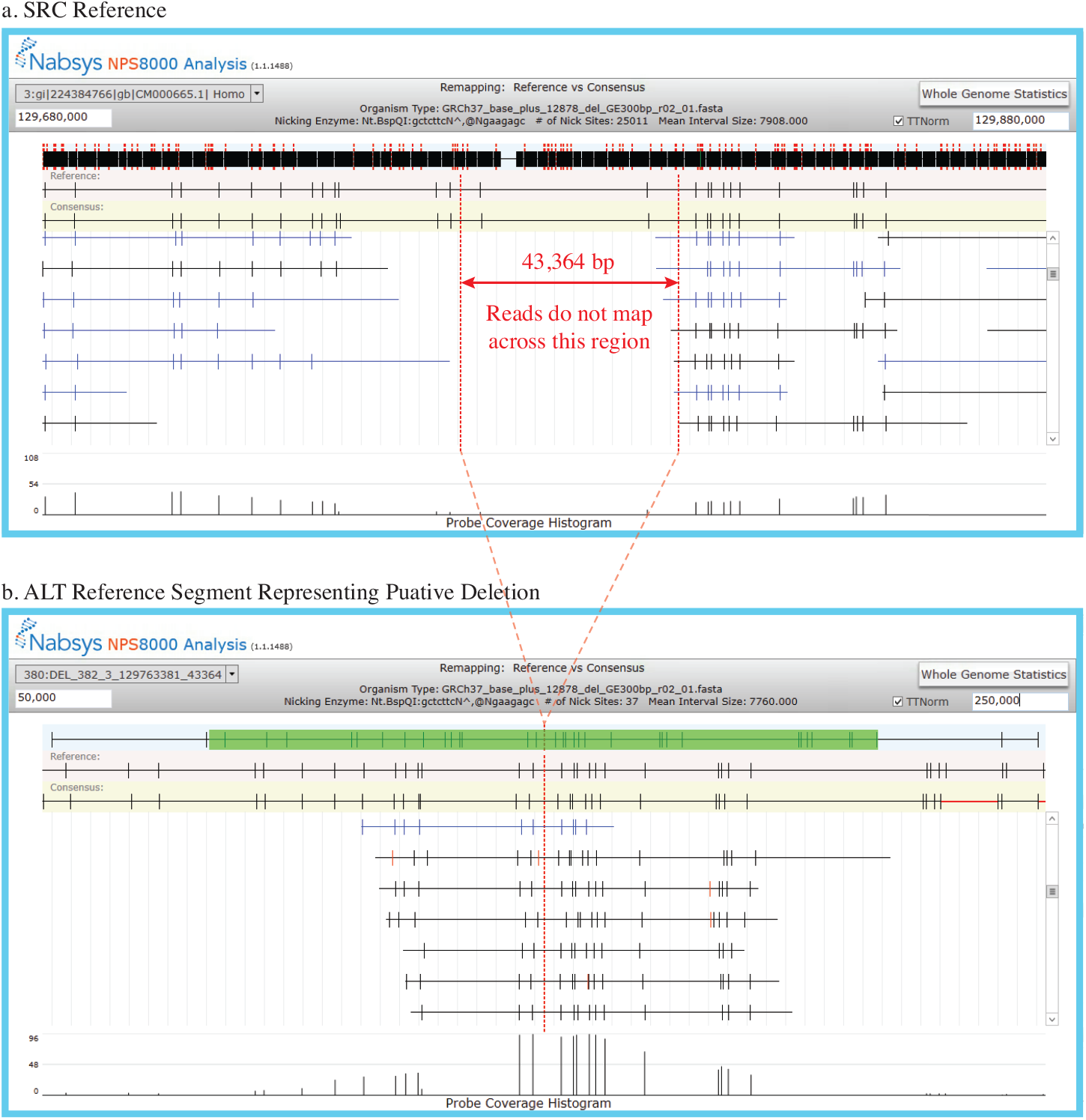
Software screen images of one homozygous deletion. Images show placement of reads on a) SRC reference and b) ALT reference segment. Only a small fraction of the reads are shown in each image and the vertical scroll bar may be used to view all mapped reads. The two red lines in a) indicate the putative breakpoints for the deletion on the SRC reference. The single red line in b) corresponds to the fusion point in the ALT reference segment.

### SVM Design and Training

The mapping method described above produces a wealth of information that can be used to inform a sophisticated variant calling approach. The *SV-Verify* software package uses a set of SVMs^23–26^ trained with high-confidence examples, to determine the likelihood that a deletion, as stated in a hypothesis, is supported by the single-molecule mapping data. The SVMs were trained with high-confidence examples of true positive deletions and true negative deletions. The resulting model, when presented with new data and a putative deletion hypothesis, outputs the posterior probability of the hypothesis being true.

The workflow used for SVM training is described in Figure 3. Separate SVMs were designed for different classes of deletion. Putative deletions were first classified based on deletion size and whether the deletion removes a probe site from the reference map. A separate SVM was trained for each class. The 4 classes were: 1) deletions that remove at least one probe site from the reference, 2) deletions 300–499 bp in size, 3) deletions 500–999 bp in size, and 4) deletions ≥ 1000 bp in size. Optimal performance was obtained using 2 sets of SVMs (8 total SVMs). For each putative deletion, we examined mapping coverage in the region of the putative deletion and used one set of SVMs when there was sufficient coverage at a stringent mapping threshold (p98) and another when there was insufficient coverage at p98, but sufficient coverage at a less stringent mapping threshold (p90). Insufficient local coverage at p90 resulted in a no-call.

**Figure 3.**
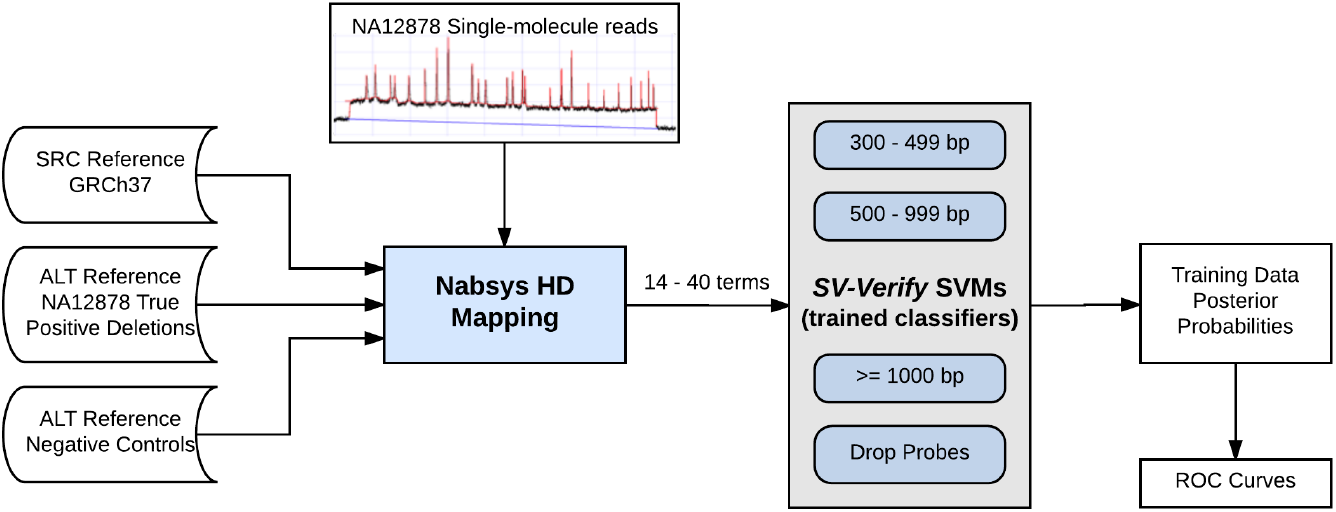
*SV-Verify* training block diagram. Data inputs for the Nabsys *HD-Mapping* software package are shown. Mapping outputs a large number of statistical parameters defining the interval size determination for every interval in the reference. The statistical parameters were used to train the SVMs.

Training proceeded by selecting an appropriate kernel^24,25^ for each SVM, selecting meaningful features from mapped single-molecule data, and applying cross-validation to avoid overfitting. Data were collected on the Nabsys platform to generate 99-fold average coverage of human genome NA12878. As indicated in Figure 4, the single-molecule reads were mapped to three different reference configurations: 1) the SRC reference alone, 2) the SRC reference plus an aggregate ALT reference that contains all putative deletions to be tested, and 3) the ALT reference alone.

**Figure 4.**
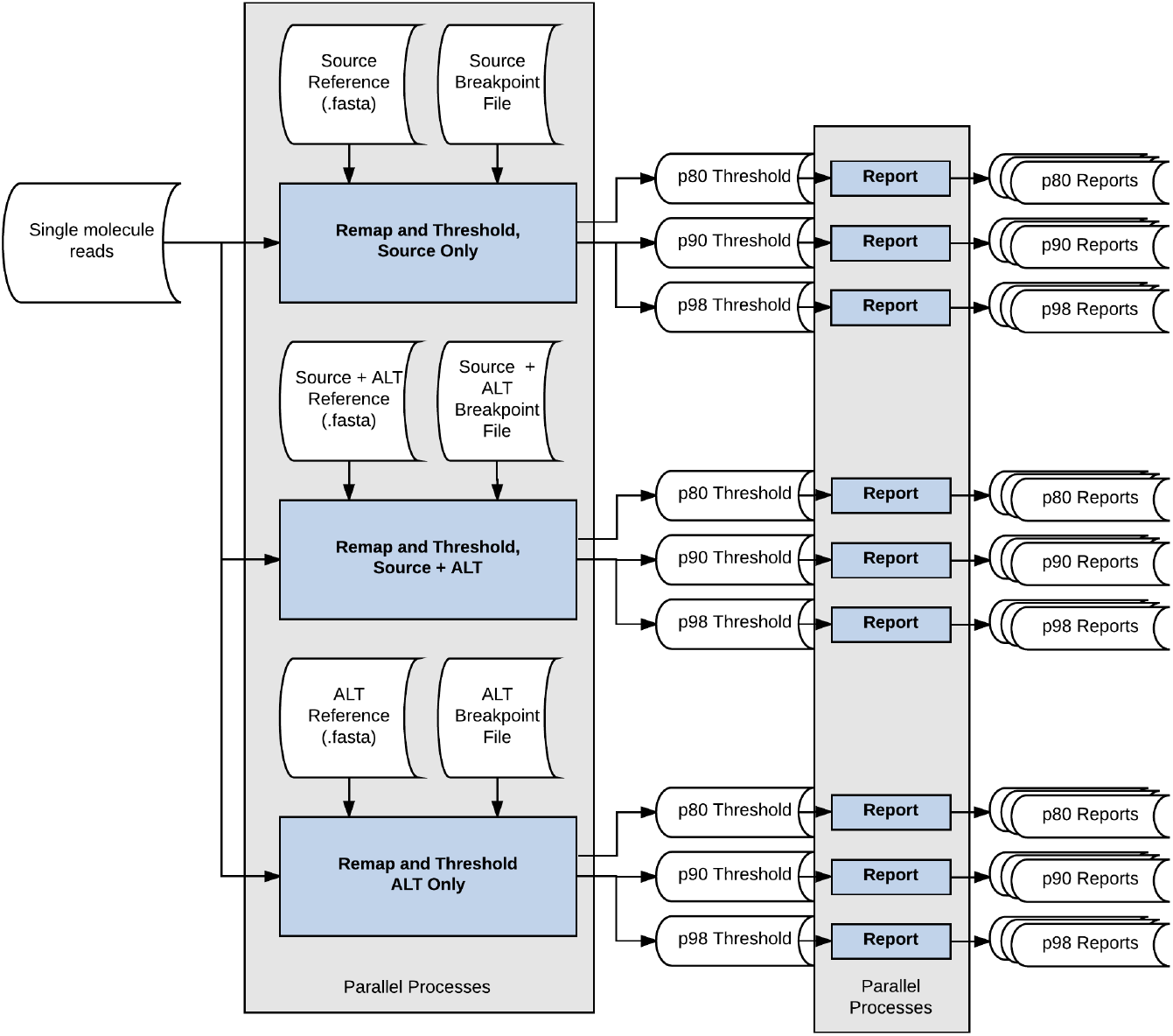
*SV-Verify* mapping detail. Single-molecule reads are the input for three parallel mapping processes each at three thresholds (p80, p90, and p98) utilizing the SRC reference alone, the SRC and ALT references together, and the ALT reference alone.

The results were evaluated at three mapping thresholds, p80 (least stringent), p90, and p98 (most stringent), with all 8 SVMs using features from all mapping thresholds. All mapping thresholds provided mapping statistics that informed all of the SVMs while the p90 and p98 results were used to determine the appropriate SVM for each putative deletion. Statistics regarding each interval in the generated SRC and ALT consensus maps, such as the first through fourth moments, were recorded and used to generate features for each SVM. For each putative deletion, we also gathered statistics related to the number of molecules mapping across SRC reference breakpoints and the corresponding ALT reference fusion point. All counts were normalized to the average coverage obtained when mapping to only the SRC reference.

### True Positive Training Data Set

While the most important requirement for true positive training data is that the vast majority of the training observations labeled as true positive must be high-confidence, it is not essential that all observations be correctly labeled. A small percentage of incorrectly labeled observations will not have a significant impact on the training.^27^ In this case, the set of true positive deletions also must be representative of the putative deletions encountered in other genomes, both in size and zygosity. We limited our true positive training set to deletions in NA12878 identified by the GIAB consortium that are greater than 300 bp and that were supported by four or more technologies. Although our consensus interval size resolution is better than 100 bp, we chose to be more conservative for our initial work and only considered putative deletions greater than 300 bp. We were cognizant that using the NA12878 high-confidence, breakpoint-resolved deletions had the potential to introduce a training bias because they are not necessarily representative of all deletions. The bias introduced by using only breakpoint resolved deletions could potentially reduce sensitivity for complex deletions, while maintaining specificity. The number of true positive deletions used to train each of the four classes of SVM is shown in Table 1.

**Table 1.** Number of deletions in the NA12878 true positive training set in each SVM class.

| Putative Deletion Size (bp) | Number of True Positive Deletions |
| --- | --- |
| 300 - 499 | 1030 |
| 500 - 999 | 205 |
| $\geq 1000$ | 387 |
| Drop Probes | 217 |

### True Negative Training Data Set

The true negative deletions are a collection of likely false hypotheses. Two true negative sets, NC01 and NC02, were each created using the following method. For every true positive deletion, we created one true negative deletion of the same size. We placed the true negative deletion at a random location in the genome, requiring that the new location’s interval size match that of the corresponding true positive interval size within 10%. We also required that mapping coverage at the new location be within 10% of the corresponding true positive mapping coverage. This last requirement is a proxy for similarity of interval sizes in the local neighborhood of the asserted deletion. Creating a true negative set in this manner permitted us to use features such as putative deletion size, interval size, and local coverage without introducing bias.

To improve specificity, for the “≥ 1,000 bp” and “drop probes” SVMs, we created an additional true negative set, NC11, consisting of randomly sized deletions, located in random genome loci. We created 1,000 deletions that deleted probe sites from the SRC reference sized between 2,000 and 50,000 bp. We created 1,000 deletions that did not delete probe sites from the SRC reference, sized between 1,100 and 10,000 bp. In all cases, we required local mapping coverage to be at least 25% of the average mapping coverage, and separated each deletion by at least 50 reference intervals. For SVMs trained using NC11, putative deletion size, interval size and local coverage could not be used as features because the difference in distributions of deletion size, interval size, and local coverage between the true positive set and the true negative sets were not constrained to be the same. Differences in these distributions are artifacts of the method used to create set NC11 and do not represent generalized features.

### SVM Feature Selection

Feature selection algorithms typically fall into one of three categories: filter, wrapper and embedded methods.^28,29^ Filter methods evaluate features by characterizing intrinsic properties of the training data. They do not involve the optimization of any specific classifier method (e.g. SVM). A wrapper method utilizes one predetermined classifier, varies features, and evaluates impact on the efficacy of the classifier. Because classifier training is required for each iteration, wrapper methods are computationally expensive. Embedded methods embed feature selection directly into the classifier construction. A gradient descent method is typically used to determine feature weights, indicating the relevance of the corresponding features. Our feature selection process utilized both filter and wrapper methods.

#### Filter methods

We graphically evaluated all potential features, in groups of 2 or 3, looking for groups of features that visually separated the training data. A 3D bubble plot showing two features that separate true positive and true negative training data is shown in Figure 5. The two features being considered are plotted on the x- and y-axes. The posterior probability from one iteration of SVM development is plotted, for reference, on the z axis. A line, y = mx, separates the majority of the members of the true positive set from the members of the true negative set, indicating the efficacy of the features under consideration.

**Figure 5.**
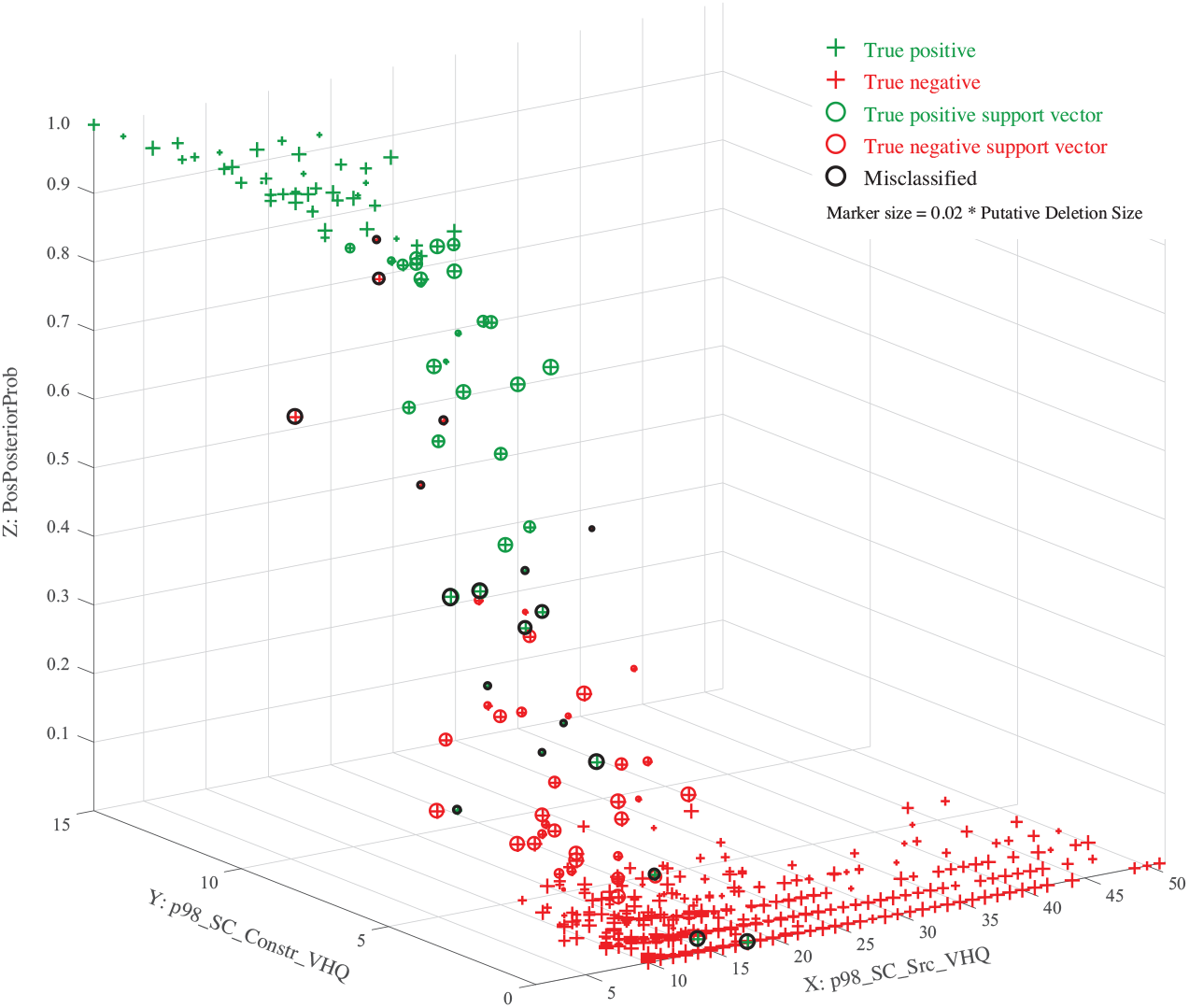
A 3D bubble plot showing two features which effectively separate training data for the “drop probes” SVM.

#### Wrapper methods

For all SVMs, other than the 300 - 499 bp SVM, we utilized a linear kernel. When using a linear kernel and normalizing features, SVM training creates a vector ***β*** whose elements may be interpreted as feature-weights. Given a matrix ***X***, containing *i* observations, each with *j* features, we created a standardized matrix, ***Xs***, with each feature normalized to mean = 0, and standard deviation = 1.

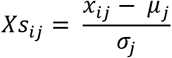

*Xs_ij_* Observation i, normalized feature j
*X_ij_* Observation *i*, feature *j*
*μ_j_* Mean of feature j
*σ_j_* Standard deviation of feature j

Using a linear kernel, function *f*(***Xs***) produces the orthogonal distance from observation ***Xs_i_*** to the hyperplane decision surface. The sign of *f*(***Xs***) indicates whether the observation is on the correct side of the hyperplane decision surface.

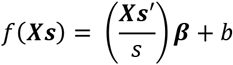

***Xs*** Normalized observations (matrix)
*s* Kernel scale
***β*** Feature weights (vector)
*b* Bias

For each linear-kernel SVM, we started by including all features, except those excluded due to known training bias. We then performed SVM training and evaluation of cross-validation accuracy, sensitivity and specificity. After training, we evaluated β, and removed features with minimal weight. We then repeated the process, iteratively removing features until there was a negative impact on training.

#### Kernel Selection

Given a training data set and the freedom to choose any SVM kernel (e.g. high-order polynomial), it is possible to achieve nearly perfect training results while producing a thoroughly inextensible, irrelevant SVM, that is unable to classify data, other than the training data. This process is known as *overfitting*. To avoid overfitting, we constrained ourselves to using kernels which were supported by visualization of the training data and required a minimum crossvalidation accuracy when performing 5-fold cross-validation^24,25^ during training.

In describing filter methods for SVM feature selection we presented an example (Figure 5) that showed two features for which a line, y = mx, separates the majority of the members of the true positive set from the members of the true negative set. In evaluating the relationships between more than 40 potential features, we typically found either no separability or linear separability. Therefore, we initially used a linear kernel for all SVMs. However, for the 300 - 499 bp SVM, we noticed a deviation from linear behavior as putative deletion size decreased and therefore evaluated a quadratic kernel. The quadratic kernel proved superior to the linear kernel for sensitivity, specificity, and 5-fold cross-validation.

## RESULTS AND DISCUSSION

### NA12878 Training Results

In order to evaluate the applicability of our trained SVMs to putative deletions in other human genomes, we performed a 5-fold cross-validation analysis during the SVM training process. A value less than 1 may either indicate mislabeling of training data (incorrect elements in the true positive or true negative training data) or less than perfect SVM performance. The results are shown in Table 2 and are consistent with expectations, confirming the efficacy of the training. SVMs utilizing the more stringent p98 mapping thresholds have better cross-validation than those using the p90 mapping thresholds. SVMs for larger putative deletions, which are easier to classify, have better cross-validation than those for smaller putative deletions.

**Table 2.**
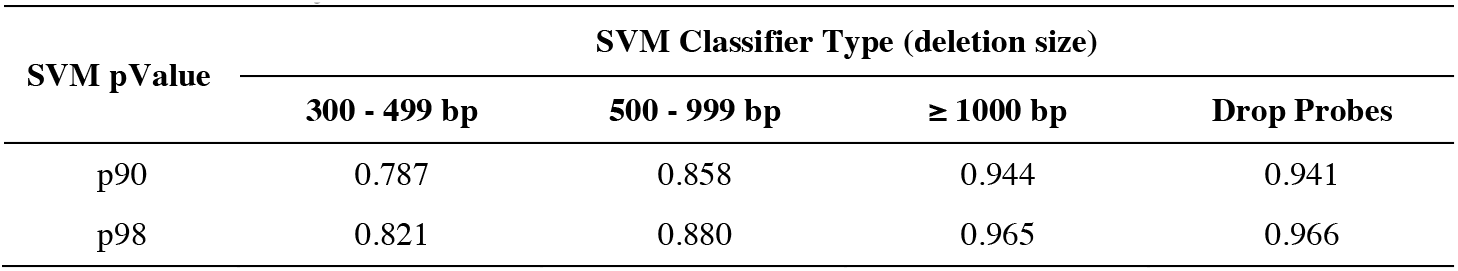
Training cross-validation accuracy.

| SVM pValue | SVM Classifier Type (deletion size) |  |  |  |
| --- | --- | --- | --- | --- |
| | 300 - 499 bp | 500 - 999 bp | $\geq 1000$ bp | Drop Probes |
| p90 | 0.787 | 0.858 | 0.944 | 0.941 |
| p98 | 0.821 | 0.880 | 0.965 | 0.966 |

We utilized the NA12878 true positive deletion call set as well as the negative control sets as training inputs for *SV-Verify* as shown in Figure 3. Results for evaluating true negative training data, using fully trained SVMs, at three specificity thresholds, *sp*90, *sp*95, and *sp*98, are shown in Table 3. For true negative data, at a given specificity threshold *sp*, the percent of evaluated deletions we expect to confirm is equal to 100 * (1 - *sp*). We confirmed 11.9%, 5.9%, and 2.1% at *sp*90, *sp*95, and *sp*98 respectively, closely matching the expected values in all cases.

**Table 3.**
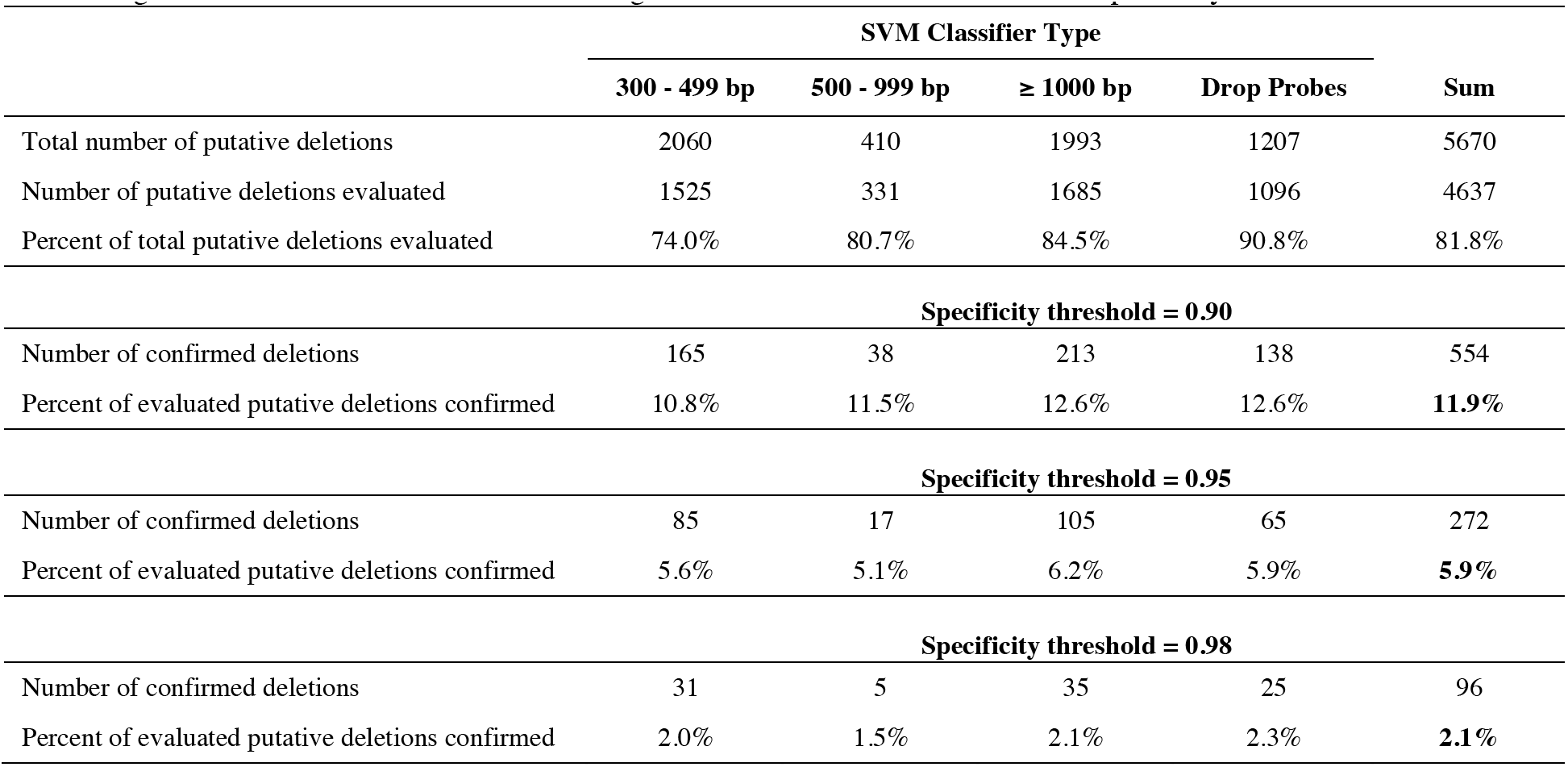
Training results of calls on the NA12878 true negative data sets for each SVM at three specificity thresholds.

Results from the evaluation of true positive training data at three different specificity thresholds are shown in Table 4. As expected, we confirm an increasing percent of the evaluated deletions as the specificity threshold is lowered across all four classifier types. We confirmed 80.8%, 72.8%, and 60.8% of true positive deletions at *sp*90, *sp*95, and *sp*98, respectively. The relatively modest decrease in sensitivity with increased specificity is highly desirable, allowing operation at high specificity with minimal impact on sensitivity.

**Table 4.** Training results of calls on the NA12878 true positive data sets for each SVM at three specificity thresholds.

|  | SVM Classifier Type |  |  |  | Sum |
| --- | --- | --- | --- | --- | --- |
| | 300 - 499 bp | 500 - 999 bp | $\geq 1000$ bp | Drop Probes | |
| Total number of putative deletions | 1030 | 205 | 387 | 217 | 1839 |
| Number of putative deletions evaluated | 729 | 142 | 294 | 182 | 1347 |
| Percent of total putative deletions evaluated | 70.8% | 69.3% | 76.0% | 83.9% | 73.2% |
| **Specificity threshold = 0.90** |  |  |  |  |  |
| Number of confirmed deletions | 548 | 110 | 264 | 167 | 1089 |
| Percent of evaluated putative deletions confirmed | 75.2% | 77.5% | 89.8% | 91.8% | **80.8%** |
| **Specificity threshold = 0.95** |  |  |  |  |  |
| Number of confirmed deletions | 476 | 96 | 258 | 151 | 981 |
| Percent of evaluated putative deletions confirmed | 65.3% | 67.6% | 87.8% | 83.0% | **72.8%** |
| **Specificity threshold = 0.98** |  |  |  |  |  |
| Number of confirmed deletions | 397 | 62 | 229 | 131 | 819 |
| Percent of evaluated putative deletions confirmed | 54.5% | 43.7% | 77.9% | 72.0% | **60.8%** |

The relationship between specificity and sensitivity is presented as a family of receiver operating characteristic (ROC) curves. A ROC curve is generated by evaluating the false positive fraction (1 - specificity) and the true positive fraction (sensitivity) as a function of posterior probability threshold. ROC curves for all eight SVMs are shown in Figure 6. On each ROC curve, the dashed orange diagonal line indicates expected performance for a random classifier and the dark green line represents perfect classification. As expected, the p98 family of SVMs has better classification performance than the corresponding p90 SVMs. All SVMs exhibit excellent specificity while retaining high sensitivity.

**Figure 6.**
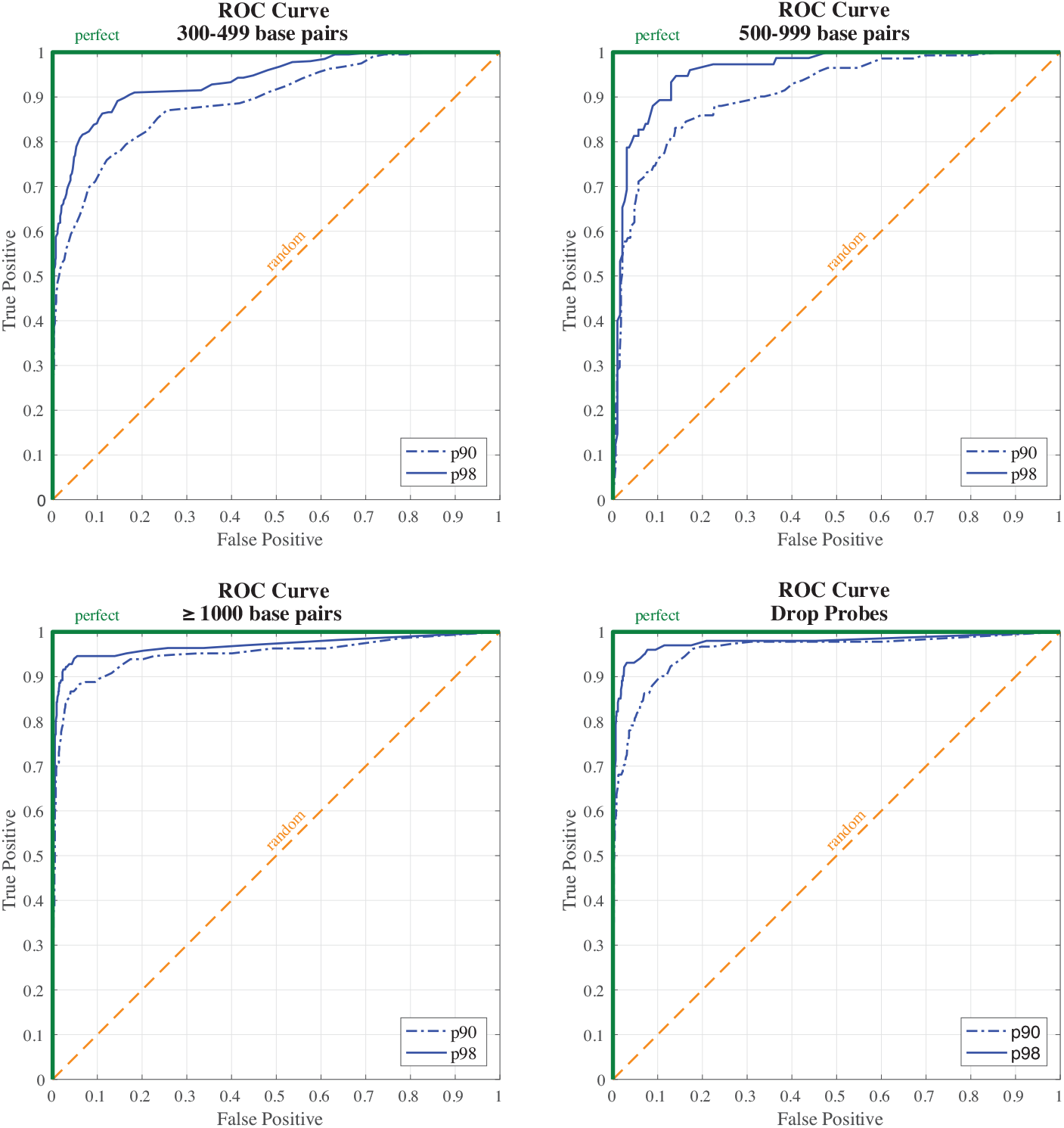
ROC curves showing the trade-off between sensitivity and specificity for each SVM at two mapping thresholds.

### Deletion verification in NA24385

In order to evaluate the performance of *SV-Verify*, we selected a putative deletion call set derived from a different human genome, NA24385. As with NA12878, the NA24385 calls were produced from multiple technologies by the GIAB consortium. While the NA12878 deletion call set contains only breakpoint-resolved calls, the NA24385 set is far less refined with > 9000 putative deletion calls ≥ 300 bp with varying degrees of support. GIAB partitioned this set into putative deletions confirmed by two or more technologies and those asserted by a single technology. We utilized *SV-Verify* to independently evaluate each of these putative deletion sets. The application of *SV-Verify* employs the ROC curves generated during training to output a specificity of detection for each putative deletion from the calculated posterior probability. The workflow for the NA24385 analysis is shown in Figure 7.

**Figure 7.**
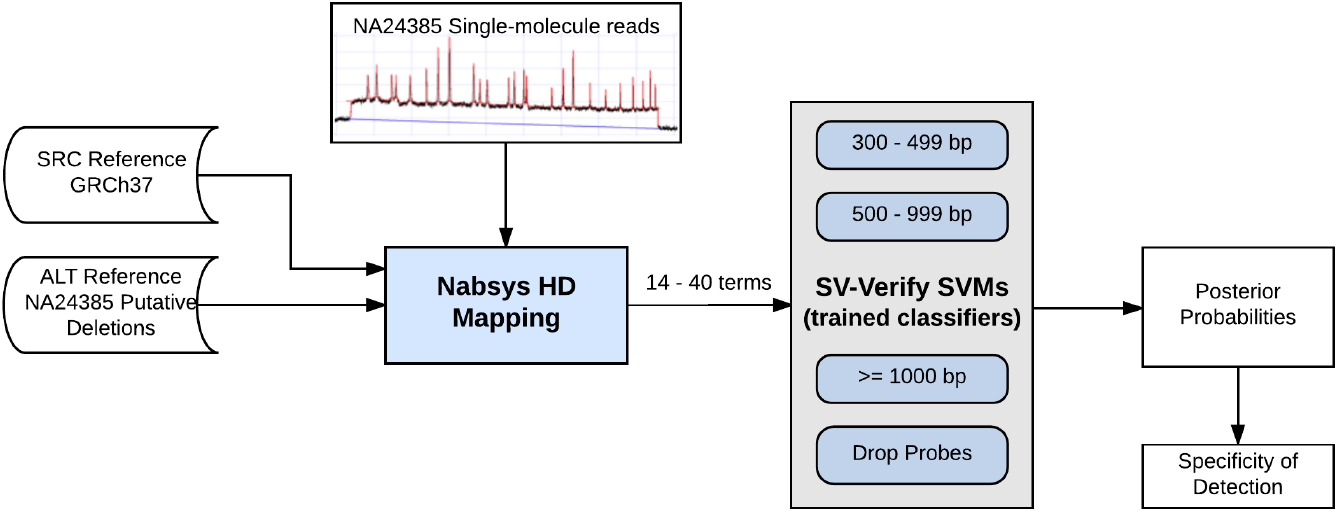
Block diagram for the application of *SV-Verify* to evaluate putative deletions in NA24385.

The results obtained by applying *SV-Verify* to the set of putative deletions asserted by only one technology (the union call set, excluding calls supported by multiple technologies) are shown in Table 5. Results obtained by applying *SV-Verify* to the set of putative deletions asserted by two or more technologies are shown in Table 6. As expected, our confirmation rate was higher for putative deletions asserted by two or more technologies. For example, at sp90, we confirmed 35.1% of evaluated deletions called by one technology versus 75.0% of evaluated deletions asserted by two or more technologies.

**Table 5.**
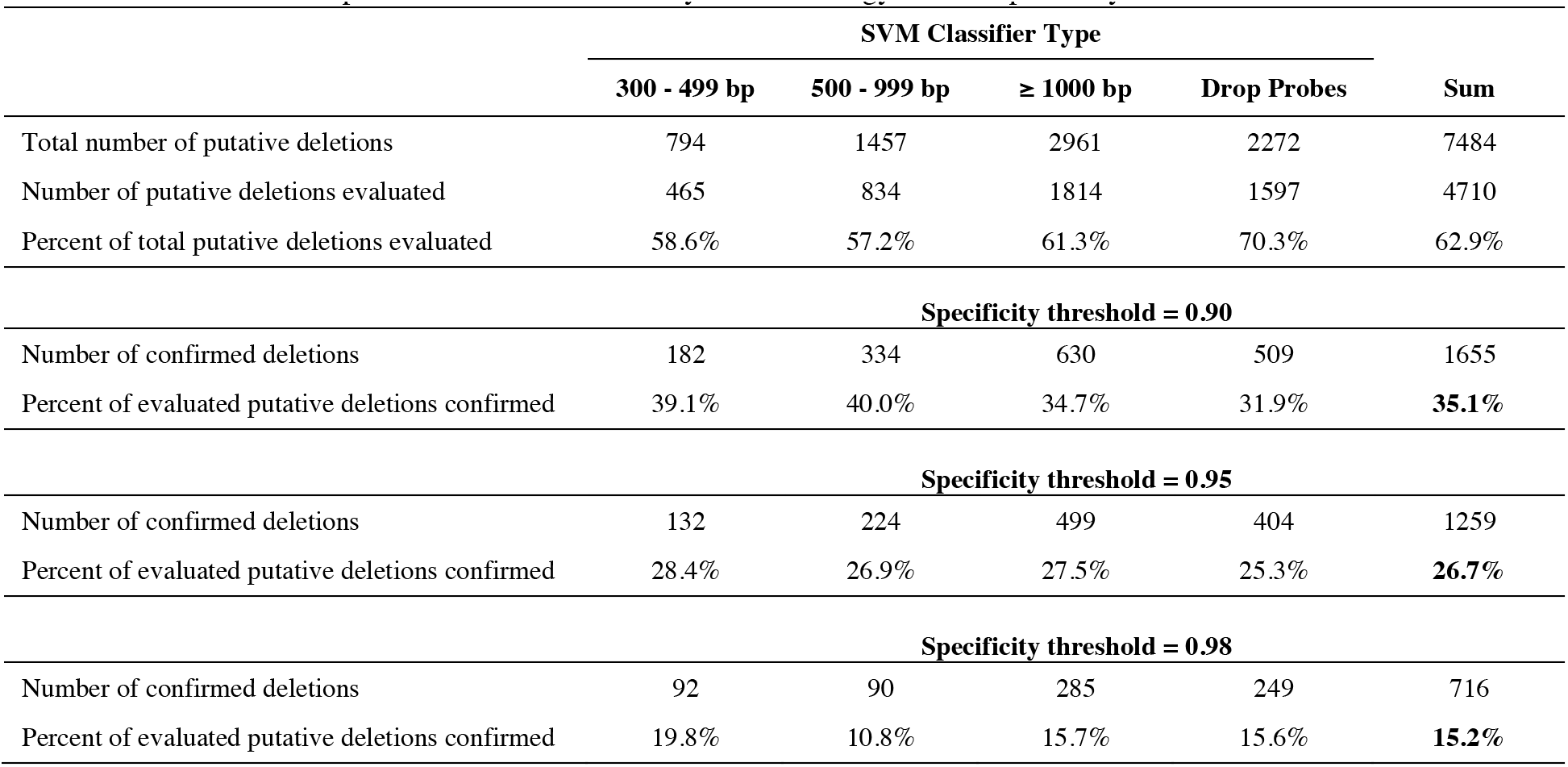
Evaluation of NA24385 putative deletions asserted by one technology at three specificity thresholds.

**Table 6.**
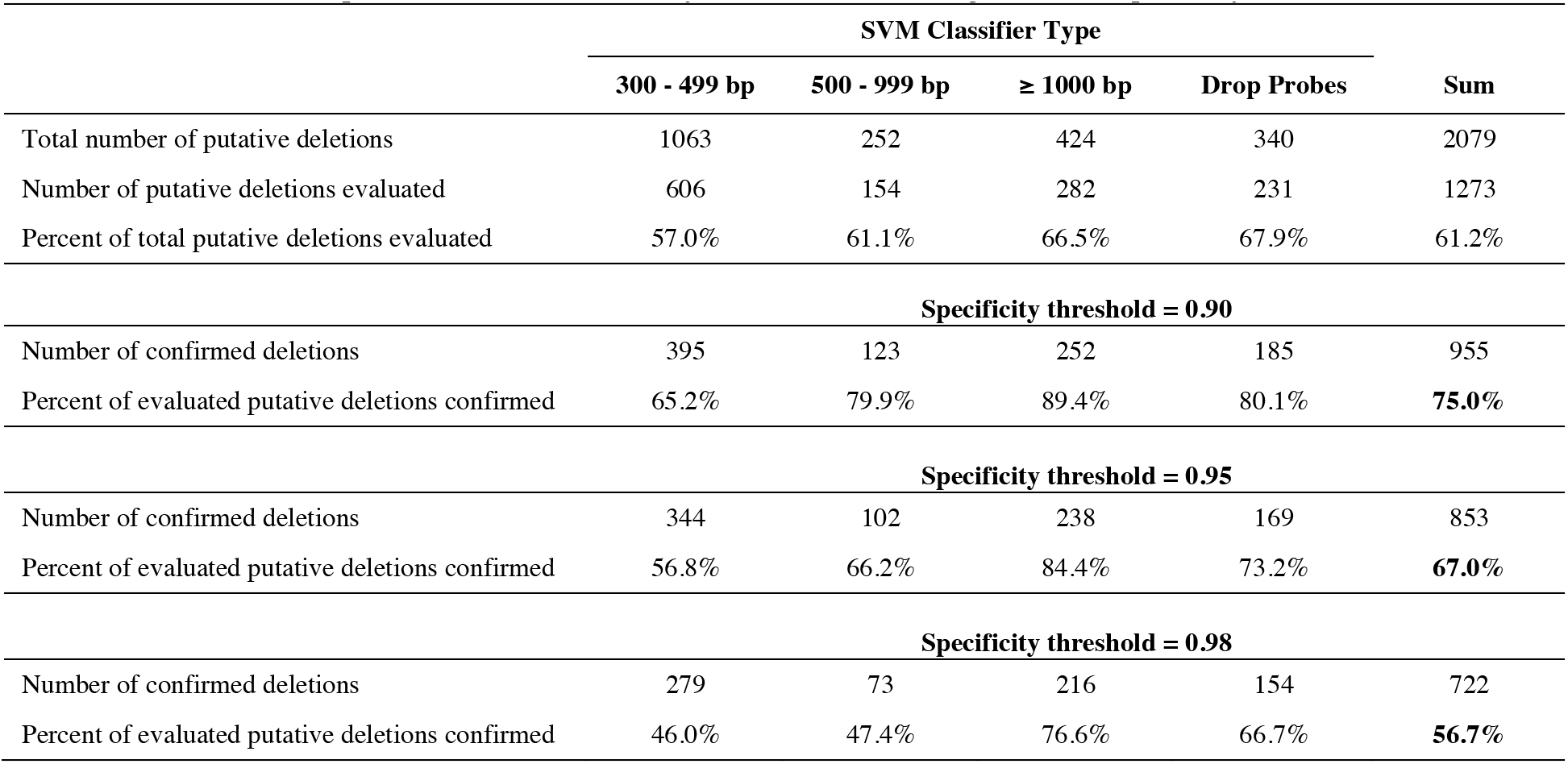
Evaluation of NA24385 putative deletions asserted by two or more technologies at three specificity thresholds.

The GIAB consortium has historically quantified the correctness of a putative deletion based on the number of independent technologies asserting the same deletion. As mentioned previously, the NA12878 high-confidence deletions were all asserted by at least four technologies (4 Tech). In order to further evaluate *SV-Verify*, we explored our NA24385 deletion calls with respect to the number of technologies that asserted the deletion. The posterior probability distribution resulting from the evaluation of all 5,983 putative deletions considered by *SV-Verify* is shown in Figure 8a. Of the evaluated putative deletions, 43% had a posterior probability < 0.1 indicating they were not supported by the data. This is not surprising, given that 78.3% of the total putative deletions were supported by only one technology. The posterior probability distributions grouped by the number of technologies asserting the same putative deletion are shown in Figure 8b. *SV-Verify* confirmed a large percentage of 4 Tech putative deletions (47% with posterior probability > 0.9) and refuted a large percentage of 1 Tech putative deletions (50% with posterior probability < 0.1). As indicated by the tailed ends of the distributions, *SV-Verify* is able to clearly discriminate between accurate calls (enriched in 4 Tech set) and inaccurate calls (prevalent in 1 Tech set).

**Figure 8.**
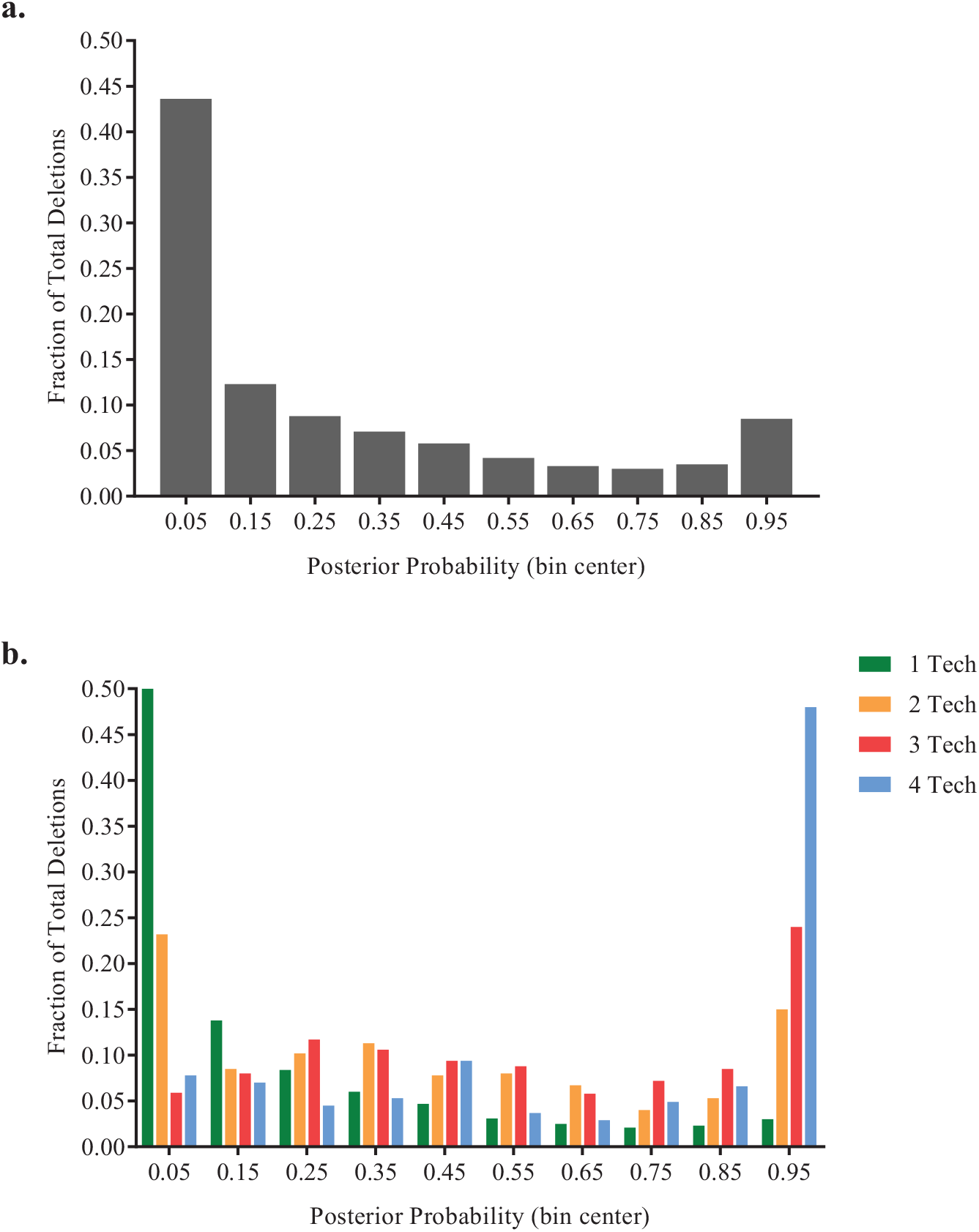
Posterior probability distributions for NA24385 putative deletion calls. The posterior probability distribution of a) all evaluated putative deletion calls or b) putative deletion calls filtered based on support by 1 (n = 4443), 2 (n = 600), 3 (n = 691), or 4 (n = 244) technologies.

When determining whether a putative deletion is confirmed, it is essential to do so at a stated specificity threshold. To convert a posterior probability, output by an SVM, to a specificity we projected the posterior probability from the SVM onto the appropriate ROC curve obtained from training (Figure 6). The percentage of evaluated deletions confirmed by *SV-Verify* at sp90 is shown in Figure 9. Results are grouped by putative deletion size and the number of technologies asserting the putative deletion. We do not present a 4 Tech result for the 300 - 499 bp range, as there were a statistically insignificant number putative deletions asserted by 4 technologies. In this size range, one of the 4 technologies, Bionano, asserted a total of 20 putative deletions, 3 of which comprise the 4 Tech intersection. The results shown in Figure 9 demonstrate that *SV-Verify* had good sensitivity across all size ranges, including 300 - 499 bp. As expected, as the number of technologies asserting a deletion increases (indicating a higher likelihood of hypothesis correctness), *SV-Verify* confirmed an increasing percentage of evaluated deletions.

**Figure 9.**
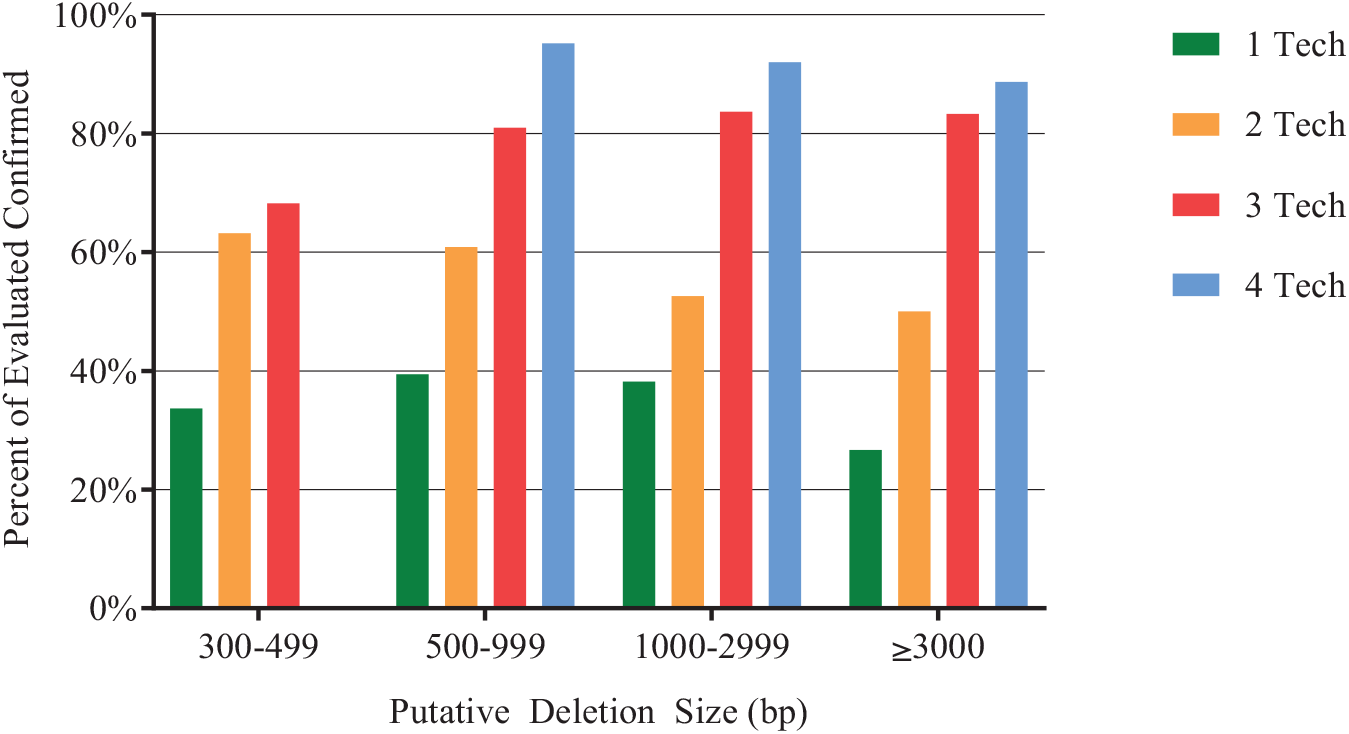
Sensitivity of NA24385 putative deletion calls at *sp*90. The percent of deletions confirmed as a function of the number of supporting technologies is plotted for different deletion sizes.

The portions of putative deletions confirmed by *SV-Verify* at specificity thresholds of *sp*90 and *sp*98 are shown in Figure 10. Each pie wedge represents the putative deletions asserted by 1, 2, 3, or 4 technologies. The area of each wedge is proportional to the number of putative deletions asserted by the specified number of technologies. In each wedge, the innermost section represents putative deletions confirmed at *sp*98. The central ring represents the putative deletions confirmed at *sp*90 and the outer ring represents putative deletions not confirmed at *sp*90. As the number of technologies increased, the percentage confirmed at any specificity threshold increased, consistent with the notion that putative deletions asserted by a larger number of technologies are more likely to be correct. Also, as the number of technologies increased, the difference between the percentage confirmed at *sp*98 and the percentage confirmed at *sp*90 decreased. For high-confidence deletions (4 Tech), the sensitivity penalty for using a high specificity threshold is minimal.

**Figure 10.**
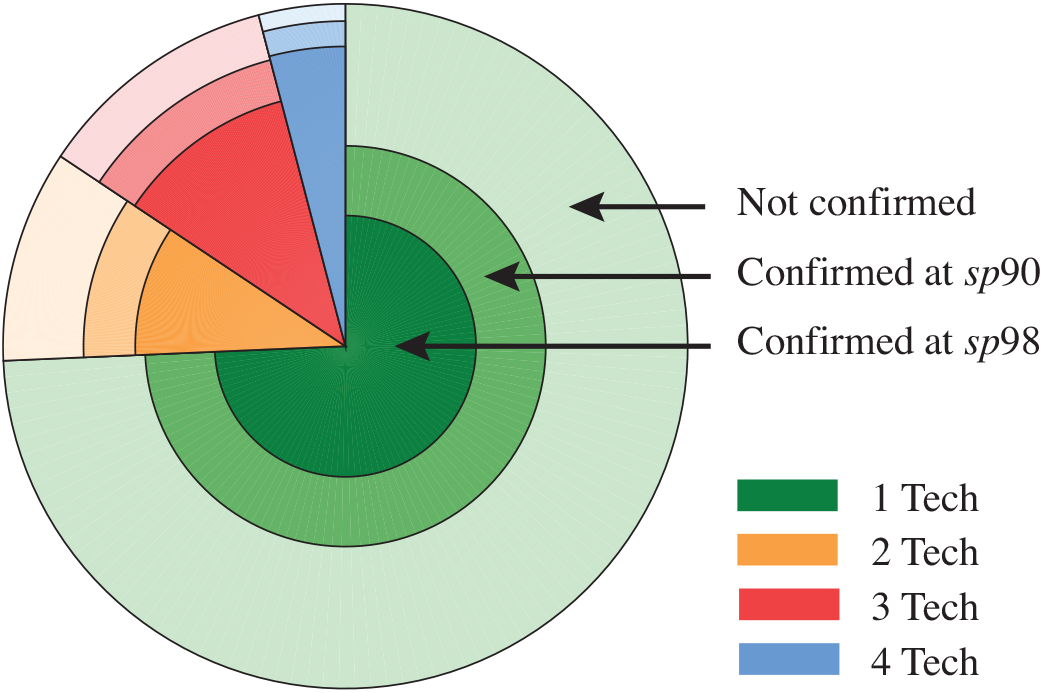
Detection sensitivity vs. number of technologies asserting the same putative deletion. The central ring indicates putative deletions confirmed at *sp*98, the middle ring indicates putative deletions confirmed at *sp*90, and the outer ring indicates unconfirmed putative deletions. Colors indicate support by 1, 2, 3, or 4 technologies. Area is proportional to the number of deletions.

The percentages of evaluated putative deletion calls made using different underlying technologies that were confirmed or refuted by *SV-Verify* in each size range are shown in Figure 11, with the corresponding numbers of calls indicated above each bar. It is important to note that the technology specifies the underlying data type, and that for each technology, there may be several data sets which were subsequently analyzed with a variety of callers. A specificity threshold sp90 was used to confirm and a posterior probability ≤ 0.1 was used to refute a putative deletion call. These threshold choices were based on ROC curves (Figure 6) and the posterior probability distribution (Figure 8). Calls that were neither confirmed nor refuted fell outside the stringent thresholds used and may represent inaccurately defined deletion hypotheses or more complex variants.

**Figure 11.**
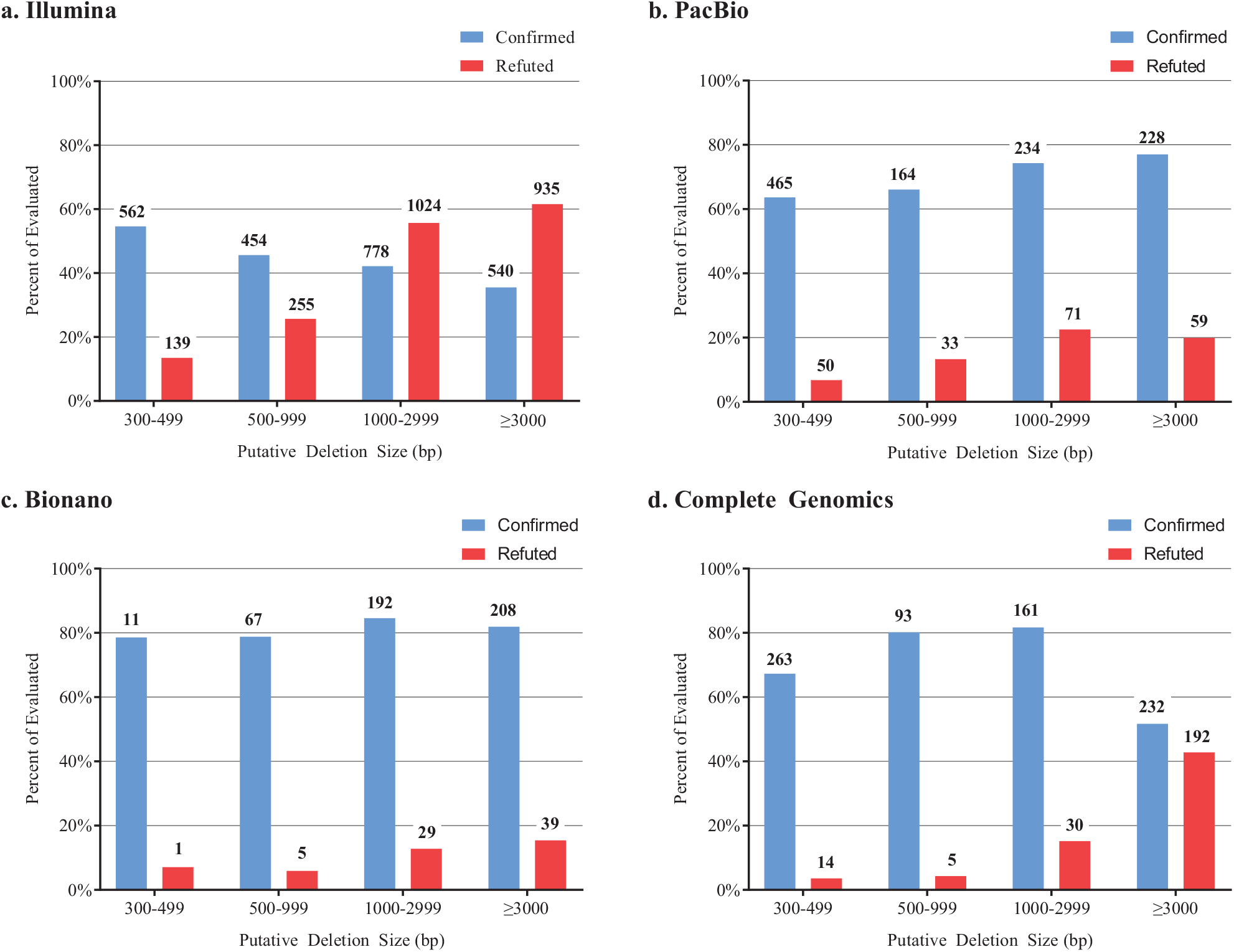
Putative deletions calls either confirmed or refuted by *SV-Verify* separated by technology. The percent of evaluated deletion calls in each size range confirmed at *sp*90 (blue bar) or refuted at ≤ 0.1 posterior probability (red bar) with data generated using a) Illumina, b) PacBio, c) Bionano, or d) Complete Genomics. Number above each bar indicates the number of deletion calls.

Putative deletions were asserted by 9 distinct callers using Illumina data sets, spanning the entire size range. These callers are tuned independently and achieve different degrees of sensitivity and specificity for different deletion sizes. The percentage of putative deletion calls confirmed decreased as a function of deletion size, while the percentage of refuted calls increased with increasing deletion size (Figure 11a). This result is likely due to the short read length associated with the underlying data. In contrast, *SV-Verify* confirmed a more consistent percentage of deletions across all size ranges from the 6 callers utilizing PacBio or the two callers using Bionano data (Figures 11b, c). Deletion calls made using Complete Genomics data by three callers were likewise similarly confirmed in the smaller size ranges, but showed a significantly higher percentage of refuted calls in the ≥ 3000 bp size range (Figure 11d). Overall these results affirm that technologies providing longer range information generate more accurate calls across a wide range of deletion sizes. In addition, these results indicate that no single technology has the sensitivity and specificity to generate a complete and accurate set of high-confidence deletion calls even when multiple callers are used.

## CONCLUSION

We have described the development of *SV-Verify*, a robust method for the verification of putative deletions called by a variety of different algorithms and underlying data types. SVMs were trained using a high-confidence deletion call set from the human genome NA12878, a powerful and unique resource produced by the GIAB consortium. For training, high-confidence NA12878 deletions served as true positives and a set of true negatives were created by asserting deletions in random areas of the genome. Single-molecule data from the Nabsys *HD-Mapping* platform have a resolution well below the limit of data from optical mapping platforms. As a result, *SV-Verify* is capable of verifying deletions as small as 300 bp and with no upper limit on size. This is demonstrated by the excellent sensitivity versus specificity characteristics displayed by ROC curves for each of the SVMs. The *SV-Verify* software package combined this effective SVM training with high-resolution single molecule *HD-Mapping* and enabled high throughput, fully automated evaluation of thousands of putative deletion calls from the NA24385 genome.

The Nabsys platform capabilities allow a seamless extension of high-confidence deletion calls to any length scale when used in conjunction with a single NGS technology, eliminating putative deletions that would not be substantiated by additional technologies. As additional high-confidence deletion data become available, *SV-Verify* will be refined to provide even higher performance for deletion calling. The methods described here are readily extensible to other types of structural variants, such as insertions, large inversions, translocations, and complex variants. Moreover, further development will enable discovery of SVs not revealed by current technologies and analytical approaches.

## ACKNOWLEDGEMENTS

We thank Justin Zook, Ph.D. and Marc Salit, Ph.D., of the NIST Genome-Scale Measurement Group for welcoming our participation in the GIAB consortium and integrating our results with those provided by other consortium members. We thank Eliezer Upfal, Ph.D., Professor of Computer Science, Brown University for his machine learning expertise and review of this manuscript.

